# Biodiversity trends are stronger in marine than terrestrial assemblages

**DOI:** 10.1101/457424

**Authors:** Shane Blowes, Sarah Supp, Laura Antão, Amanda Bates, Helge Bruelheide, Jon Chase, Faye Moyes, Anne Magurran, Brian McGill, Isla Myers-Smith, Marten Winter, Anne Bjorkman, Diana Bowler, Jarrett E.K. Byrnes, Andrew Gonzalez, Jes Hines, Forest Isbell, Holly Jones, Laetitia M. Navarro, Patrick Thompson, Mark Vellend, Conor Waldock, Maria Dornelas

**Affiliations:** German Centre for Integrative Biodiversity Research (iDiv), Halle-Jena-Leipzig, Germany; Data Analytics Program, Denison University, Granville, OH, USA; Centre for Biological Diversity, School of Biology, University of St. Andrews, St. Andrews, UK; Department of Biology and CESAM, Universidade de Aveiro, Aveiro, Portugal; Department of Ocean Sciences, Memorial University of Newfoundland, Newfoundland, Canada; Martin Luther University Halle-Wittenberg, Institute of Biology / Geobotany and Botanical Garden, Halle (Saale), Germany; Martin Luther University Halle-Wittenberg, Institute of Computer Science, Halle (Saale), Germany; School of Biology and Ecology, University of Maine, Orono, ME, USA; School of GeoSciences, University of Edinburgh, Edinburgh, UK; Senckenberg Gesellschaft für Naturforschung, Biodiversity and Climate Research Centre (BiK-F), Frankfurt am Main, Germany; Department of Biology, University of Massachusetts Boston, Boston, MA, USA; Department of Biology, McGill University, Montreal, Qc, Canada; Leipzig University, Institute of Biology, Leipzig, Germany; Department of Ecology, Evolution, and Behavior, University of Minnesota, St. Paul, MN, USA; Department of Biological Sciences and Institute for the Study of the Environment, Sustainability, and Energy, Northern Illinois University, DeKalb, IL, USA; Department of Zoology, University of British Columbia, Vancouver, BC, Canada; Département de biologie, Université de Sherbrooke, Sherbrooke, QC, Canada; Ocean and Earth Science, National Oceanography Centre, University of Southampton, Southampton, UK & Life Sciences, Natural History Museum, Cromwell Road, London, UK.

## Abstract

Human activities have fundamentally altered biodiversity. Extinction rates are elevated and model projections suggest drastic biodiversity declines. Yet, observed temporal trends in recent decades are highly variable, despite consistent change in species composition. Here, we uncover clear spatial patterns within this variation. We estimated trends in the richness and composition of assemblages in over 50,000 time-series, to provide the most comprehensive assessment of temporal change in biodiversity across the planet to date. The strongest, most consistent pattern shows compositional change dominated by species turnover, with marine taxa experiencing up to fourfold the variation in rates of change of terrestrial taxa. Richness change ranged from no change to richness gains or losses of ~10% per year, with tropical marine biomes experiencing the most extreme changes. Earth is undergoing a process of spatial reorganisation of species and, while few areas are unaffected, biodiversity change is consistently strongest in the oceans.

## Main Text

Biodiversity is changing rapidly throughout the Anthropocene. Against a background of elevated extinction rates ^1,2^ local biodiversity change results from multiple interacting drivers that influence the abundance and distribution of species. Different regions of the globe are projected to experience different trends in biodiversity change, particularly due to variation in the strength of drivers such as land use intensity^3^ and climate change^4^. There are widespread changes in the identities of species that live in any one location (species composition), whereas shifts in the numbers of species (species richness) show a mixed pattern with increasing, decreasing, or static trends^8–8^. However, the spatial distribution of the locations most affected is unknown. Here, we map biodiversity change, as species richness and composition, to establish whether there are systematic trends in the biogeography of biodiversity change. Our analysis compares marine and terrestrial realms, as well as different biomes, and latitudinal bands examined as polar, temperate, and tropical regions of the globe.

Biodiversity and its change is unevenly distributed on the planet^9,10^, and unevenly sampled^14–14^. Detecting geographic variation in biodiversity trends will inform conservation prioritisation and improve estimates of global biodiversity change. Moreover, quantifying this spatial distribution will help refine hypotheses about the drivers of biodiversity change. The spatial distribution of drivers of biodiversity change is heterogeneous^15,16^ and fundamentally differs between the marine and terrestrial realms^17^. Specifically, there is more spatial overlap between climate change and other drivers of change in the marine realm than in the terrestrial realm^17^. Understanding the biogeography of biodiversity change across realms is essential for reliable forecasting future change and its consequences.

Quantifying biogeographic patterns of biodiversity change will allow us to assess the ongoing spatial re-organisation of species. This reorganisation is being driven by climate change driven range shifts^4,18^, altered species abundance due to land-use change^3^ and widespread species introductions^19^. Local or regional richness will decline when species losses exceed species gains, for example, in areas where land-use intensity is high^3^ and/or when range sizes contract^20^. Conversely, local or regional richness will increase when species gains exceed losses, occurring, for example, in places where species are introduced ^23–23^ and where ranges expand ^24,25^, or when species are favoured by land-use change^26^. Combinations of different anthropogenic drivers can lead to increases or decreases, depending on the magnitude of each driver^27^.

Here, we quantified biogeographic variation in patterns of change in both species richness and composition from a compilation of over 50,000 local assemblage time series (ranging from 2 to 97 years; mean = 5.5 years) across the globe. We use the BioTIME database, which is currently the largest collation of assemblage time series (332 studies analysed^28^; plus 26 other studies; see Table S1 in Dornelas et al. 2018; Extended Data Fig. 1). The null expectation is that overall species richness should remain largely stable even when compositional turnover is high, due to species gains and losses at local scales being approximately balanced and widespread community regulation^31–31^. Results from a number of analyses are consistent with this expectation^5–8,29^. Yet there is scope for non-random spatial patterns within the distribution of local species gains, losses and compositional change. Here, we quantify variation in biodiversity change trends among realms, biomes, latitudinal bands and taxa.

## Biodiversity trends across the globe

To examine biogeographical patterns in biodiversity change we estimated trends in richness and composition change using hierarchical linear models. After standardising for spatial extent and sampling effort of each time-series, we quantified biogeographic variation by examining departures from overall trends for 48 biomes and 9 taxonomic-habitat groupings (hereafter referred to as *taxa*; amphibians, benthos, birds, fish, invertebrates, mammals, marine invertebrates/plants, plants, and multiple taxa) that were nested within biomes (resulting in 105 biome-taxa combinations [table S1]). We also examined the robustness of our biome-taxa models by fitting a second set of simpler hierarchical models, where taxonomic groups were nested within latitudinal bands (i.e., polar, temperate, tropical) for each realm (marine, terrestrial, freshwater; [table S2] *see supporting information for details*).

Our results show that variation in biodiversity change is greater in the marine versus the terrestrial and freshwater realms. The overall trend in richness change was not statistically distinguishable from zero (Fig. 1), neither globally, nor at the biome level. This result was robust to sensitivity analyses regarding time series duration and start year (Extended Data Figs. 4 to 7). However, variation in departures from the overall trend were almost four times greater among marine biomes (median biome departure range: −0.0014 -- 0.0017, *n* = 33; Fig. 1a) compared with terrestrial and freshwater biomes (−0.0006 -- 0.0002, *n* = 15; Fig. 1b). Thus, taxa in marine biomes frequently represented extremes at both ends of the range of observed change in species richness - negative trends of approximately 10% species loss per year and positive trends approaching 15% species gains per year (Fig. 2a). At the taxa-level, this volatility in marine species richness change meant that a higher proportion of biome-taxa combinations were undergoing richness changes that differed from zero with 95% probability (36/78) compared with terrestrial and freshwater taxa (7/27).

**Figure 1.**
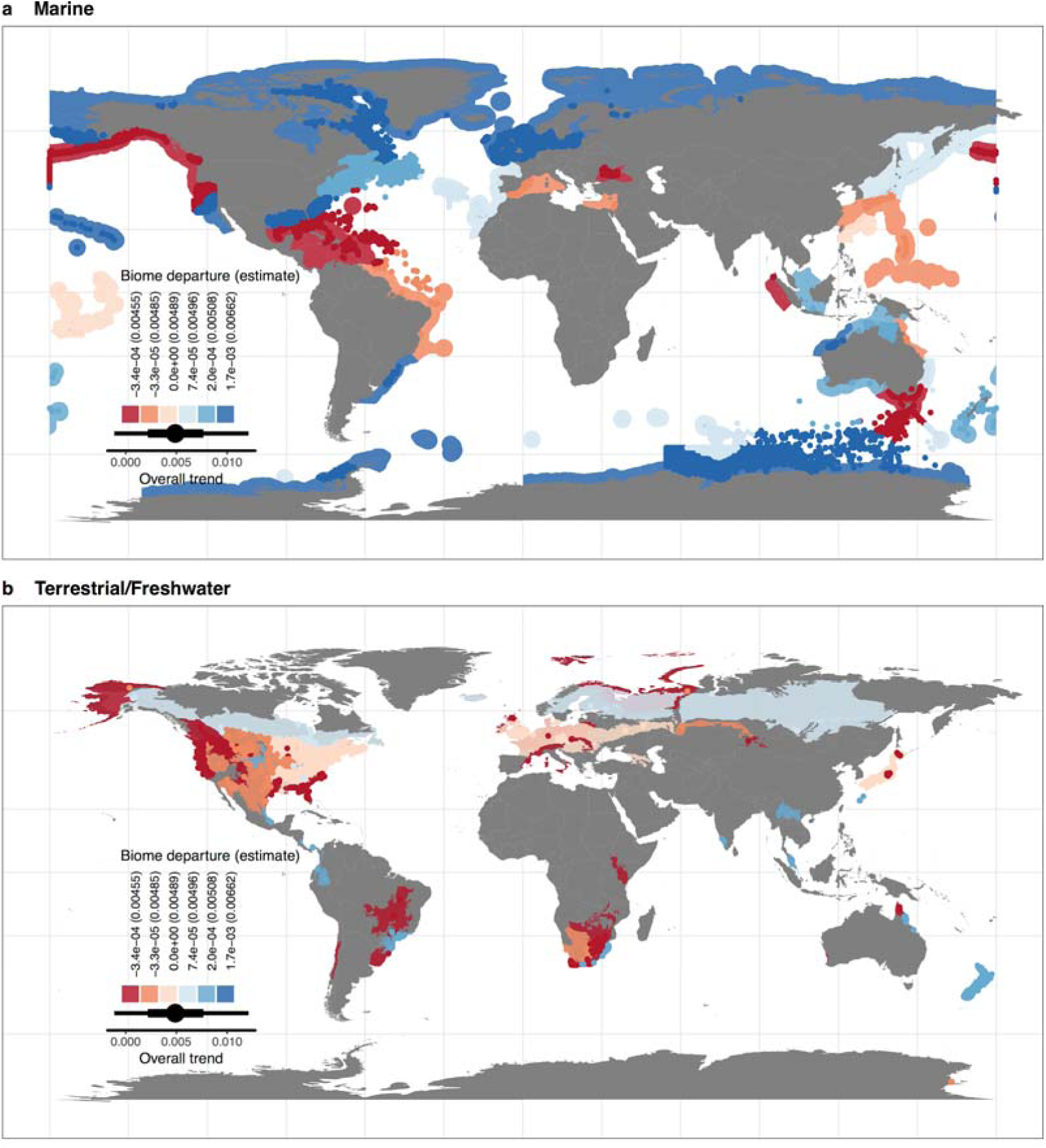
The overall trend in assemblage species richness change across biomes in the marine and terrestrial realm does not differ from zero; bar depicts 50% (thick) and 90% (thin) credible intervals. Shading represents positive (blue) and negative (red) departures from the overall trend (0.005) for each biome; numbers in legend denote the departure and the biome-level (overall + departure) estimate in brackets. 90% credible intervals for all biome level estimates overlap zero. **a**, Marine biomes (n = 33) show both positive and negative departures from the overall trend, with more negative departures in the tropics, whereas there are no latitudinal trends in **b**, Terrestrial (n = 10) and freshwater (n = 5) biomes. Marine biomes show stronger variation in richness outcomes; particularly visible in the dark blue coastal polar regions and the cluster of dark red biomes in the Caribbean and southeast coast of Australia.

Richness trends varied substantially across taxa within biomes, and this variation was spatially structured, with the most pronounced trends found within tropical latitudes. Negative richness trends (i.e. slope estimates with 90% credible intervals that do not include zero) were present for taxa in five out of twelve marine tropical biomes, with positive trends present in three (Fig. 2a). These results were consistent with our simpler hierarchical model that showed overall declines within tropical latitudes for marine taxa (Fig. 2b). Locations where species losses outweigh gains could be driven by range contractions, or by the loss of more specialised or thermally restricted species as climate change, land use or seascape change, and other anthropogenic drivers affect tropical habitats^32,33^. Geographic gaps from terrestrial tropical systems remain in our assemblage time series data, precluding direct comparison between the realms (see also Extended Data Fig. 1). The high rates of change we observed in the marine tropics are consistent with predictions that tropical species will be relatively sensitive to extreme heat events, because they are closer to their physiological limits, resulting in biodiversity loss^34^. However, overexploitation, pollution, and other threats are likely also contributing to biodiversity change.

**Figure 2.**
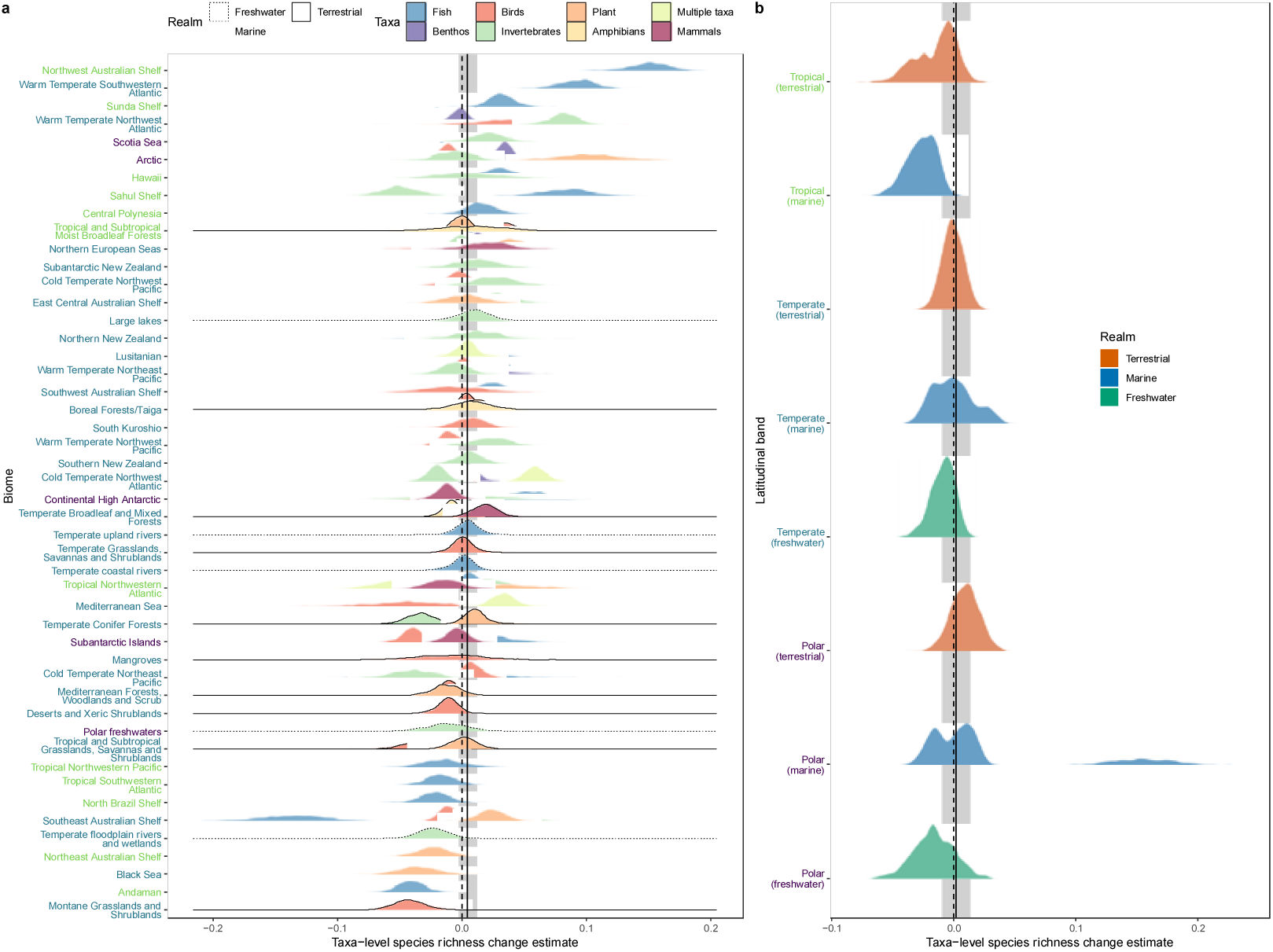
The magnitude of species richness change is more variable among taxa in marine biomes than taxa in terrestrial or freshwater biomes. The overall trend in assemblage richness change (solid vertical bar) does not differ from zero (grey shading depicts the 90% credible interval) for both **a**, the biome-taxa model, and **b**, the simpler realm-climate-taxa model. Density ridges in **a** represent the posterior distribution for the slope coefficients of each taxa-level coefficient (within each a biome); line type and fill refer to the realm and taxon, respectively. Colour of biome label denotes latitudinal band (light green tropical, blue green temperate and purple polar as shown in **b**). Density ridges in **b** represent the posterior distributions of the slope coefficients for all taxa within a given combination of realm and latitudinal band (climate) estimated with the realm-climate-taxa model.

To examine changes in species composition, we partitioned total Jaccard dissimilarity, calculated as the dissimilarity between the initial and each subsequent year of a time series, into the additive components of turnover and nestedness^35^. These trends describe directional compositional change relative to the initial assemblage, and the decomposition examines whether changes in community composition were due to the original species in assemblages being replaced by new species (turnover), or if assemblages were becoming smaller subsets of themselves or growing to include new species alongside the original species (nestedness). Overall, we found rates of *turnover* were positive and much greater (0.025; 90% credible interval: 0.021-0.029; Fig. 3) than the rate of change in *nestedness* (0.005; 0.004-0.006; Extended Data Fig. 2). This means that compositional change was dominated by species replacement within assemblages, with approximately 25% of species within assemblages being replaced per decade. Marine biomes showed both positive and negative departures from the overall trend (i.e., depending on the biome, more and less compositional change over time than the global average; Fig. 3a). In contrast, the terrestrial biomes showed mostly negative departures from the global average (Fig. 3b), often in highly developed regions of the globe (e.g., Northeast US, Europe, Japan). Similar to our finding for species richness change, variation in rates of turnover were more than 1.5 times greater in marine biomes (Fig. 3) and 2.5 times greater among marine taxa when compared to their terrestrial and freshwater counterparts (Fig. 4). Taxa in terrestrial and freshwater biomes represented 9 of the 10 lowest rates of turnover, whereas, 9 of the 10 highest rates of turnover were marine taxa (Fig. 4a). Fish in marine tropical biomes represented both ends of the spectrum, from among the lowest turnover rates (Tropical Southwestern Atlantic) to the highest observed turnover rate (Northwest Australian Shelf).

**Figure 3.**
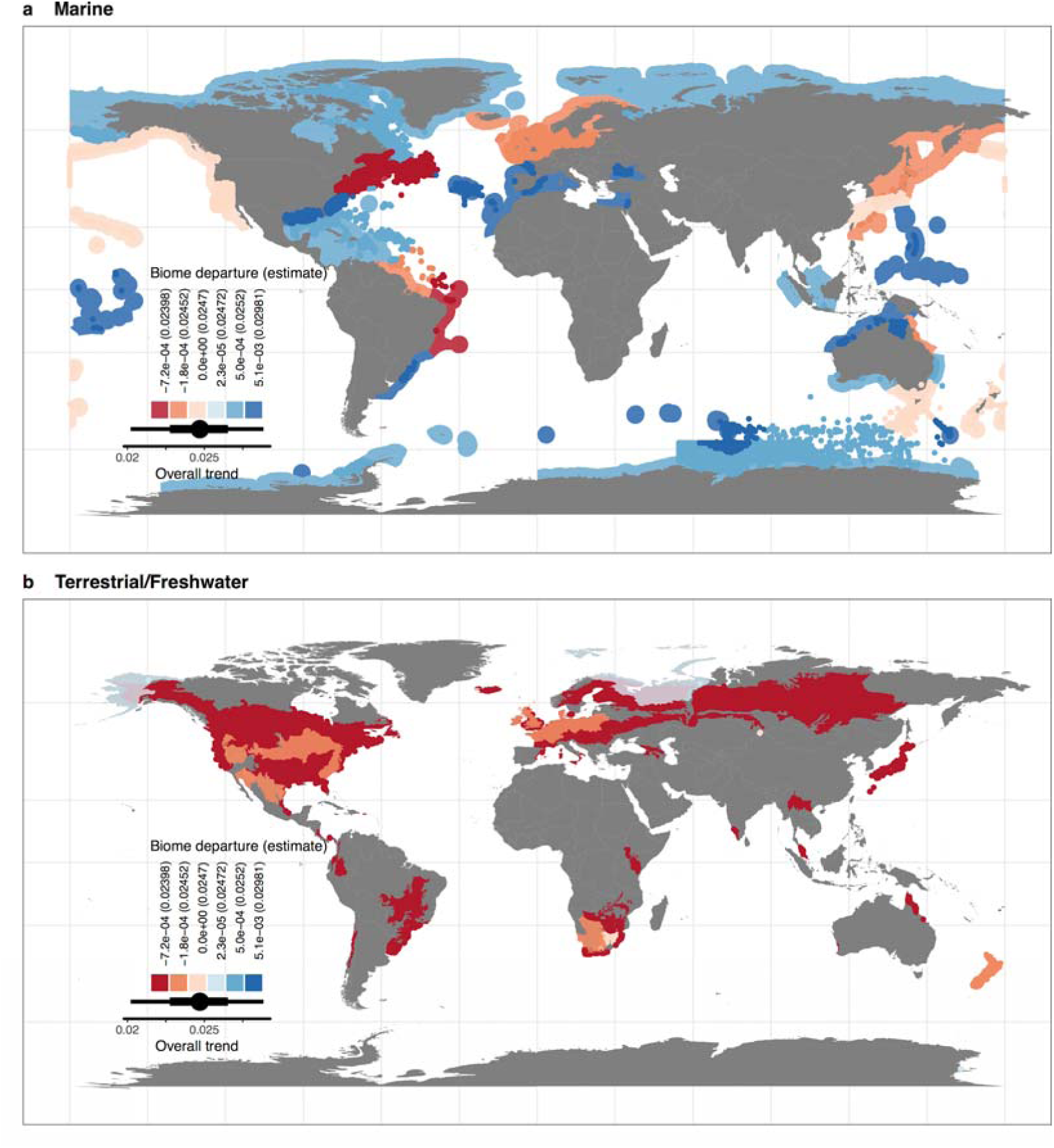
Assemblages across the globe are experiencing high rates of species replacement (turnover component of Jaccard dissimilarity). **a**, Rates of new species replacing original species have both positive and negative departures from the overall trend in marine biomes, whereas **b**, terrestrial and freshwater biomes have mostly slower rates of turnover than the overall trend (red shading). Numbers in legend denote the departure and the biome-level (overall + departure) estimate in brackets.

**Figure 4.**
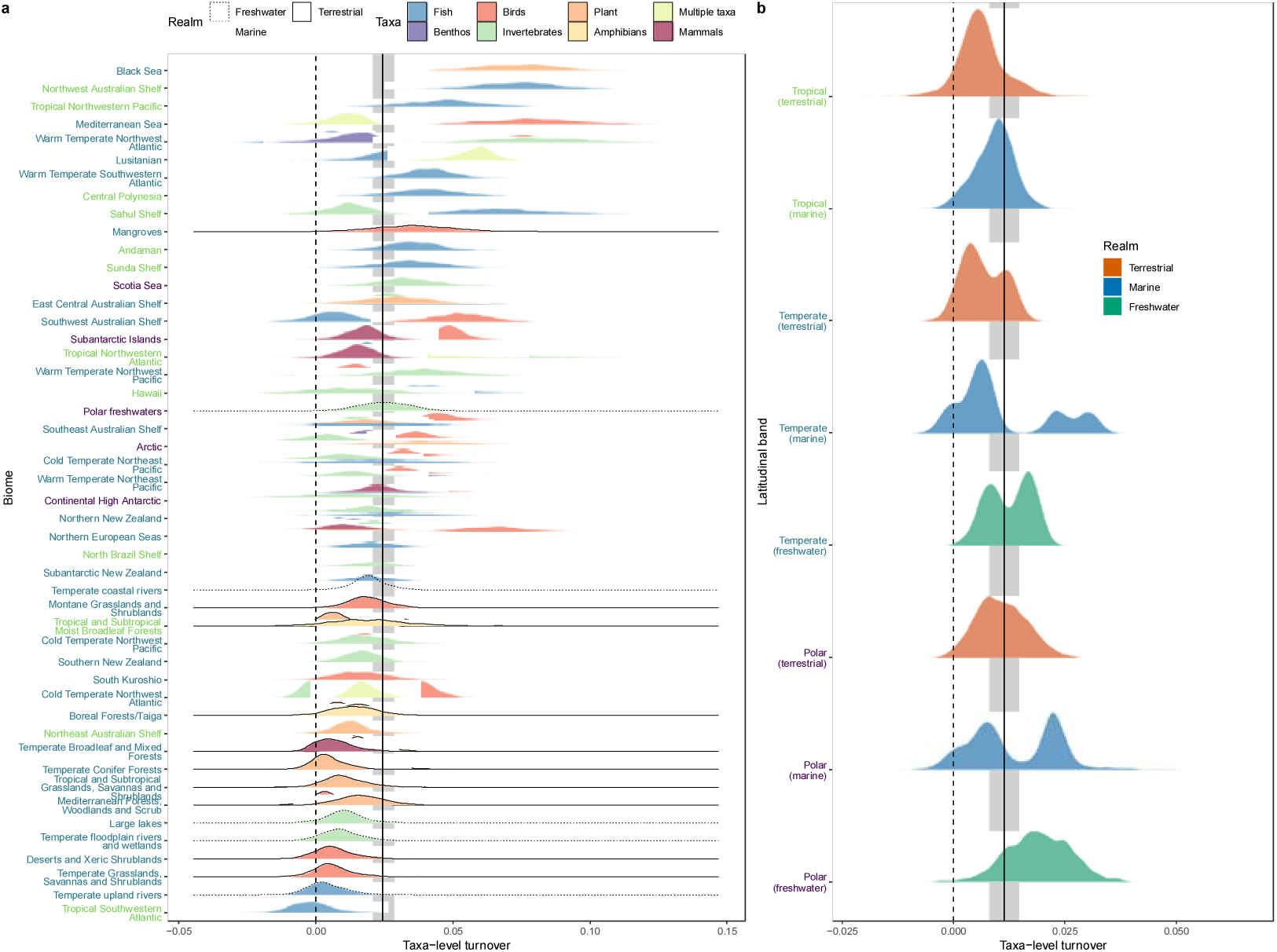
The magnitude and total variation of composition change (represented by the turnover component of Jaccard dissimilarity) is greater among taxa in marine biomes compared to taxa in terrestrial or freshwater biomes. The overall trend in turnover change per year is greater than zero (solid black line; grey shading depicts the 90% credible interval) for both **a**, the biome-taxa model, and **b** the simpler realm-climate-taxa model. Density ridges in **a** represent the posterior distribution for each taxa-level slope coefficient (within each a biome); linetype and fill refer to the realm and taxon, respectively. Colour of biome label denotes latitudinal band (light green tropical, blue green temperate and purple polar as shown in **b**). Density ridges in **b** represent the posterior distributions of the slope coefficients for all taxa within a given combination of realm and latitudinal band (climate) estimated with the realm-climate-taxa model.

## Linking richness and composition change

To illustrate the relationship between trends in species richness and composition, we plotted the dominant component of composition change (turnover or nestedness) for each biome-taxa combination as a function of species richness change (Fig. 5a, b). When turnover is the dominant component, this relationship shows how fast new species are replacing original species, and whether or not these arrivals influence the total number of species in assemblages. We found rates of turnover change to exceed nestedness for more than 90% of biome-taxa combinations (97/105; Fig. 5b, c, d). For these taxa, approximately one-third (31/97) had both rates of turnover and species richness trends different from zero (Fig. 5 c), with a balanced distribution of 14 cases of species richness losses, and 17 with gains. When nestedness is the dominant component, this relationship shows how fast assemblages are changing to become smaller subsets of the species initially observed. Among the eight biome-taxa combinations where nestedness exceeded turnover change (8/105; Fig. 5b, c, d), three marine taxa had increasing nestedness and detectable species richness losses. Spatial patterns of biodiversity change were highly heterogeneous (Fig. 5d), with locations in close proximity experiencing distinct trends in composition and richness change. For example, within the cold temperate northwest Atlantic marine biome (Fig. 5d, Extended Data Figure 3; encompassing the Gulf of Maine, Gulf of St. Lawrence, and south Newfoundland), the trend for invertebrate assemblages to become a smaller subset of themselves was in close proximity to five other groupings of taxa where turnover was the dominant component of composition change (benthos, birds, fish, marine invertebrates/plants and multiple taxa), among which there were trends for both species losses (e.g. birds) and gains (e.g. multiple taxa).

**Figure 5.**
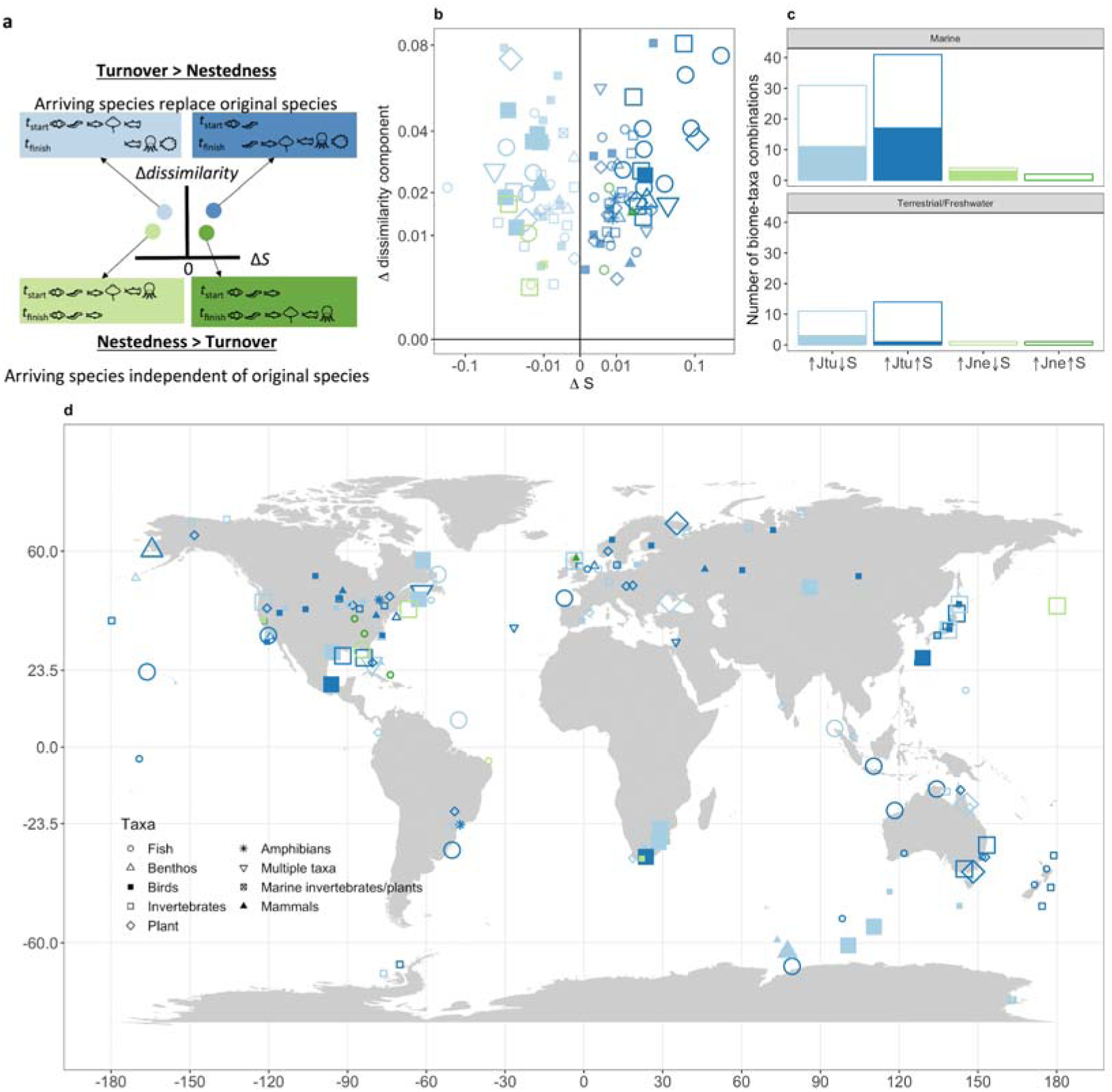
Conceptual and empirical relationships between changes in species richness and composition. **a**, Conceptual model relating the turnover and nestedness components of species composition change (Δ dissimilarity) to changes in species richness (ΔS). When the turnover component is larger than the nestedness component, new species entering assemblages replace the original species (blue shaded boxes). Conversely, when the nestedness component is larger than the turnover component, some original species of the assemblage remain, and the numbers of new species entering the assemblage are largely independent of the original species (green shaded boxes). The change in species richness documents the net change in the numbers of species in the assemblage (and ignores their identity as either original or new species). **b**, Scatterplot showing the dissimilarity trend as a function of the species richness trend. Each point represents a taxa-level group (shapes denoted in panel **d** legend), and larger points indicate that both composition and richness trends differed from zero with 95% probability; CIs not shown and axes transformed for clarity. **c**, The number of taxa for each combination of change in species richness and species composition (measured as either the turnover, Jtu, or nestedness, Jne, component of Jaccard dissimilarity). Filled sections of each bar represent taxa where both species richness (S) and composition change (Jtu or Jne) differ from zero. **d**, Map showing the location of each biome-taxa combination. Point colours and sizes follow from panel **b**.

## Discussion

Compositional change dominated by species turnover is the most striking and most prevalent pattern in biodiversity patterns across the globe. For the majority of taxa in biomes across the planet for which we have data, there was considerable replacement of species through time with no associated species richness changes. The consistent pattern of species replacement is likely underpinned by a diverse suite of drivers. Such reorganization is consistent with regulatory mechanisms for species richness. Community regulation in species richness is widespread^30^, and may be driven by shared resource availability^29^ or by the continuous replacement of transient species^36,37^. Contemporary pressures such as introduced species^23–23^, replacement of localised specialists by widespread generalists, or range shifts in response to environmental change ^18,38,39^ may also help explain our finding of widespread composition change and variable richness change with an overall trend not distinguishable from zero.

Rates of species richness change and turnover were higher and more variable in the marine realm. Detecting this difference between realms generates hypotheses about the differences among realms in both drivers of biodiversity change and biotic responses to these drivers. The spatial distributions of anthropogenic drivers of change differ among realms^17^ and different driver combinations have different effects on biodiversity change^27^. In addition, marine organisms may be more likely to respond to some drivers such as climate change^34^ and marine assemblages may be able to turnover more rapidly due to higher connectivity in the marine realm^40^.

Amid widespread variation in biodiversity trends, we detect a signal of tropical marine regions having a distribution of trends more skewed towards richness extremes and high turnover. Particularly concerning are the two tropical marine biomes that show both negative trends in species richness and higher than average turnover of fish assemblages: Andaman and Tropical Northwestern Pacific. The tropics, which harbour the majority of the planet’s biological diversity, are generally considered to be the most threatened regions of the planet^41^. Moreover, in the context of climate change, there are likely fewer species available to replace species lost in tropical zones that have entered no-analog warm temperature conditions^42,43^. If these trends are maintained, this pattern could lead to dramatic global losses of biodiversity and to the attenuation of the latitudinal diversity gradient, significantly altering the planet’s biogeography.

Here, we identify hotspots of biodiversity change: key areas that represent increasing and decreasing extremes for biodiversity trends. In addition to the marine tropics, areas like Tasmania, Alaska, and the South of South Africa stand out as regions experiencing stronger negative biodiversity change. In contrast, the North Sea and Eastern North America emerge as areas experiencing increases in biodiversity change. This spatial and taxonomic variation in biodiversity trends means that global trends of biodiversity need to be based on spatially representative data. However, and despite using the largest compilation of biodiversity time series to date, our analysis suffers from many blind-spots. Our results highlight how improving our understanding of biodiversity change will require filling the gaps by improving biodiversity monitoring and moving towards global stratified random sampling of biodiversity.

In summary, biodiversity change has strong biogeographic variation. We have identified hotspots of richness gains and losses, as well as species replacement. The marine realm emerges as having the strongest change, and the marine tropics in particular, as having a higher prevalence of richness losses. This spatial variation suggests we need to abandon a view of homogenous loss of biodiversity, as the mean of local change across the globe does not differ from zero, and is not necessarily representative of local trends. Our work suggests an urgent need to better understand ***why*** there is such geographic variation. The spatial variation described here should inform conservation prioritisation by identifying the parts of the planet changing most rapidly, as well as those that are more stable. In the field of climate science, there was a shift in wording from global warming to climate change. Similarly, our results justify a shift in focus towards recognising that biodiversity change in the Anthropocene has contrasting effects in different parts of the planet.

## Methods

### Data description and pre-processing

The BioTIME database represents the largest global effort mobilizing assemblage time series (range = 2 to 97 years; mean = 5.5 years), includes 386 studies, and currently holds over 12 million records of abundance for over 45 thousand species across plants, invertebrates, fish, birds and mammals^28^. Analyses presented in this study are based on time series of abundance data (i.e., the studies that recorded counts of the number of individuals for each species in an assemblage).

As we were interested in quantifying biodiversity change at the local scale, studies with multiple sampling locations and extents greater than 71.7 km^2^ were partitioned into 96 km^2^ grid cells (studies with extents < 71.7 km^2^ were assigned to the grid cell in which they were centered). Each cell was given a unique identifier that was the concatenation of the study ID and the cell reference number. Species were then collated within each grid cell for each year, resulting in new assemblage time series within grid cells. For all assemblages in every cell and every year, we calculated the coverage of the sample, which is a measure of sample completeness^44^ (mean = 0.95, sd = 0.11), and removed all assemblage-cell-year combinations with coverage less 0.85 (meaning that for the time series retained, there was a <15% chance of another individual sampled being a new species).

Finally, we applied sample-based rarefaction to the minimum number of samples per year for each time series. We calculated community dissimilarity using pairwise Jaccard dissimilarity and the first year as the baseline, and species richness for each year of the time series. We also calculated the turnover and nestedness components of Jaccard dissimilarity to assess if the changes in compositional diversity were driven by species replacement or changes in species richness^35,45^.

### Models of biodiversity change

We quantified biodiversity change using two complementary hierarchical models. Both models included the time series of assemblage dynamics within individual cells at the lowest level of the model, but differed in the way that these individual time series were grouped hierarchically. The biome-taxa (BT) model nested time series within ecological biomes (http://maps.tnc.org/gisdata.html ^49–49^) and groups of taxa. The realm-climate-taxa (RCT) model nested time series within a grouping covariate that was the concatenation of realm (marine, terrestrial or freshwater), climate (latitudinal bands denoting polar regions, temperate regions and the tropics) and taxa. The groupings of taxa for both models were based on the metadata of BioTIME, and included: amphibians, benthos, birds, fish, invertebrates, mammals, marine plants/invertebrates, plants, and multiple taxa for studies that measured more than one taxa group. We fit models where there were more than three cell-level times series per group, and discuss trends at the biome and taxa levels for the BT model, and for the realm-climate-taxa level for the RCT model, as the analytic technique is not well suited to describing trends at the cell-level where the data are sparse. Both models were fit with year as a population (or global) parameter, and year (i.e., the slope parameter) and the intercept were allowed to vary for each of the hierarchical levels of the models.

We quantified change in species richness and community composition (total Jaccard dissimilarity, and the turnover and nestedness components). Species richness was modelled assuming a Poisson error distribution and a log link function. This resulted in the BT model having the form:

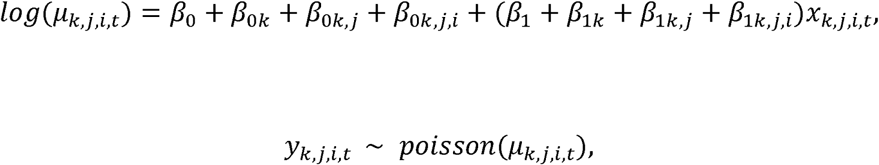

where *X_k,j,i,t_* is the time in years, *β_0_* and *β_1_* are the global intercept and slope (often termed fixed effects), *β_0k_* and *β_1k_* are the biome-level departures from *β_0_* and *β_1_* (respectively; biome-level random effects), *β_0k,j_* and *β_1k,j_* are taxa-level departures (nested within biomes) from *β_0_* and *β_1_* (taxa-level random effects), *β_0k,j,i_* and *β_1k,j,i_* are the (nested) cell-level departures from *β_0_* and *β_1_* (cell-level random effects); *y_k,j,i,t_* is the (rarefied) species richness in year *t* of the ith cell for the *j*th taxonomic group within the *k*th biome.

The species richness RCT model had the form:

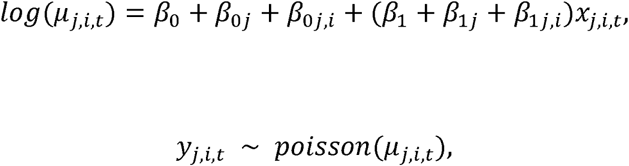

where *X_j,i,t_* is the time in years, *β_0_* and *β_1_* are the global intercept and slope (fixed effects), *β_0j_* and *β_1j_* are the departures for each realm-climate-taxa group from *β_0_* and *β_1_* (respectively; random effects), *β_0j,i_* and *β_1j,i_* are the cell-level departures from *β_0_* and *β_1_* (cell-level random effects); *Y_j,i,t_* is the (rarefied) species richness in year *t* of the ith cell, of the *j*th combination of realm-climate-taxa.

All dissimilarity metrics were modelled assuming Gaussian error and an identity link function, resulting in BT models of the form:

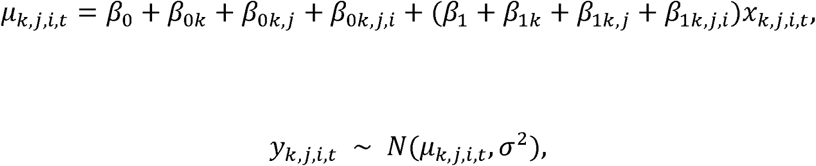

where *X_k,j,i,t_* is the time in years, *β_0_* and *β_1_* are the global intercept and slope, *β_0k_* and *β_1k_* are the biome-level departures from *β_0_* and *β_1_* (respectively), *β_0k,J_* and *β_1k,j_* are the taxa-level departures from *β_0_* and *β_1_*, *β_0,k,j,I_* and *β_1,k,j,I_* are the cell-level departures from *β_0_* and *β_1_*, and *Y_k,j,i,t_* is the value of the dissimilarity metric (total Jaccard dissimilarity, or one of the components) in year *t* of the *i*th cell, of the *j*th taxonomic group within the *k*th biome. The dissimilarity metric was set to equal zero (perfectly similarity) for the first year of each time series.

The dissimilarity RCT models had the form:

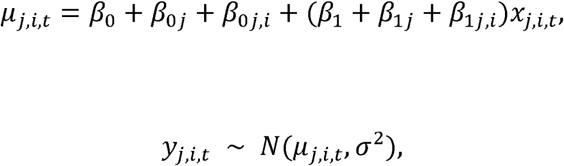

where *X_j,i,t_* is the time in years, *β_0_* and *β_1_* are the global intercept and slope, *β_0j_* and *β_1j_* are the departures for each realm-climate-taxa group from *β_0_* and *β_1_* (respectively), *β_0j,i_* and *β_1j,i_* are the cell-level departures from *β_0_* and *β_1_* and *Y_j,i,t_* is the value of the dissimilarity metric in year *t* of the *i*th cell, of the *j*th combination of realm-climate-taxa. The dissimilarity metric was set to equal zero (perfectly similarity) for the first year of each time series.

We used weakly regularising normally-distributed priors for the global intercept and slope:

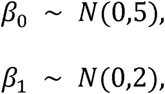

Group-level parameters were all assumed to be *N*(0, σ), and priors on the σ were the same for all models of composition (the turnover and nestedness components of Jaccard’s dissimilarity) and the RCT of species richness (i.e., as follows, with the *k* subscript dropped for the RCT models):

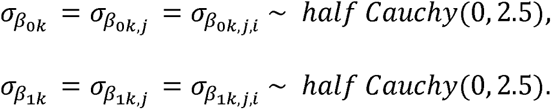

The group-level parameters of the BT species richness model were also assumed to be *N*(0, σ), but the priors were drawn from the student *t* distribution:

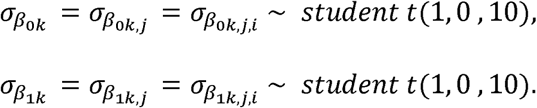

Correlations between levels of the grouping-factors (e.g., taxa with biomes) are estimated using the Cholesky decomposition (L) of the correlation matrix, with a Lewandowski-Dorota-Joe (LKJ) prior^50^, here set as:

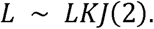

Model convergence and goodness of fit were assessed using a combination of statistics (Gelman-Rubin diagnostic^51,52^) and visual inspection of the Markov chains. All data manipulation and analysis were conducted in R (3.3.1 or greater^53^). Models were coded using the ‘brms’ package (version 1.5.1 or greater^50^), which fits models with the probabilistic programming language Stan^50^.

## Sensitivity analyses

A recurrent criticism of existing time series analyses is the lack of an appropriate baseline from which to detect change^11,54^. Obtaining baselines for all of the datasets in the BioTIME database is unrealistic, but we assessed whether the rates of change are themselves changing through time by quantifying biodiversity change for different time periods (since the 1950’s). To do this, we subset the data into three periods: 1951-1970, 1971-1990, and 1990-2010, and refit each of the models to each of these subsets (Extended Data Figs. 4-6).

We also assessed the sensitivity of our estimates of biodiversity change to the length of the time series (number of discrete years sampled), and the starting year of each time series (Extended Data Fig. 10). Additionally, we examined the estimates of change as a function of the initial assemblage species richness (i.e., the number of species in the first year of each assemblage time series; Extended Data Fig. 10).

## Code availability

Code is available on an online archive at Zenodo: (doi: https://doi.org/10.5281/zenodo.1475218)

## Data availability

The time series analysed were from 332 unique references found in the BioTIME dataset and in other studies which were used with permission. Approximately 92% (306 references) of the biodiversity studies analysed here are available as part of the published BioTIME Database^28^. The data are openly available, and can be accessed on Zenodo (https://doi.org/10.5281/zenodo.1211105) or through the BioTIME website (http://biotime.st-andrews.ac.uk/).

Dornelas, M., Antão, L.H., Moyes, F., Bates, A.E., Magurran, A.E., and BioTIME consortium *(200+ authors).* 2018. BioTIME: a database of biodiversity time-series for the Anthropocene. Global Ecology and Biogeography. 10.1111/geb.12729

The remaining 8% (26 references) of biodiversity studies analysed were used with permission. Some of these studies are published and publicly available outside of the BioTIME database, and others are available with permission from the corresponding author on reasonable request. For more details regarding the original citations, corresponding authors, and availability of these datasets, please refer to Table S1 in Dornelas et al. (2018).

## Acknowledgements

We thank all the authors who contributed their data, or gave us permission to include it in this study. Mary O’Connor helped in securing the funding for the working group and contributed to discussions and editing the manuscript drafts. This work was supported by funding to the sChange working group through sDiv, the synthesis centre of iDiv, the German Centre for Integrative Biodiversity Research Halle-Jena-Leipzig, funded by the German Research Foundation (FZT 118). Many working group members contributed to the projects in various roles across multiple meetings, and were influential in shaping the discussions and ideas that led to this paper. Those not listed as authors here include: Mary O’Connor, Juliano Sarmento Cabral, Jillian Dunic, and Robin Elahi. SB, HB, JMC, JH and MW were supported by the German Centre for Integrative Biodiversity Research (iDiv) Halle-Jena-Leipzig. SRS was supported by National Science Foundation grant 1400911. LHA was supported by Fundação para a Ciência e Tecnologia, Portugal (POPH/FSE SFRH/BD/90469/2012). MD was supported by a Leverhulme Trust Fellowship. AEM, FM and MD were supported by ERC AdG BioTIME 250189 and PoC BioCHANGE 727440.

## Author Information

These authors contributed equally and led the paper: Shane Blowes and Sarah Supp. Maria Dornelas was the senior author on the paper.

## Author Contributions

MD conceived the project. SB, SRS, and MD led the development of the project, assisted with data analysis and interpretation, and wrote the first draft of the manuscript. SB and SRS collaborated on the core data preparation and coding the analysis in R. SB designed the analytical models and prepared the figures for the manuscript. FM managed the BioTIME database, queried it for the analysis, and provided help with figures. All authors contributed to the sDiv working group that conceived this project, to key discussions that led to the design of the study, and helped with revising the paper and preparing and approving it for publication. MD and SRS secured the funding from sDiv and co-led the sChange working group (with Mary O’Connor) that initiated the project.

## Corresponding Author

Correspondence and requests for materials should be addressed to the lead authors: Shane Blowes, Sarah Supp, and Maria Dornelas.

## Supplementary Information

Further information detailing the results, including the sensitivity analysis, is available in the Supplementary Information and Extended Data files, linked to this paper.

